# ProteinFlow: a Python Library to Pre-Process Protein Structure Data for Deep Learning Applications

**DOI:** 10.1101/2023.09.25.559346

**Authors:** Elizaveta Kozlova, Arthur Valentin, Aous Khadhraoui, Daniel Nakhaee-Zadeh Gutierrez

## Abstract

Over the past few years, deep learning tools for protein design have made significant advances in the field of bioengineering, opening up new opportunities for drug discovery, disease prevention or industrial biotechnology. However, despite the growing interest and excitement surrounding these tools, progress in the field is hindered by a lack of standardized datasets for benchmarking. Most models are trained on data from the Protein Data Bank (PDB), the largest repository of experimentally determined biological macromolecular structures. But filtering and processing this data involves many hyperparameter choices that are often not harmonized across the research community. Moreover, the task of splitting protein data into training and validation subsets with minimal data leakage is not trivial and often overlooked. Here we present ProteinFlow, a computational pipeline to pre-process protein sequence and structural data for deep learning applications. The pipeline is fully configurable and allows the extraction of all levels of protein organization (primary to quaternary), allowing end-users to cater the dataset for a multitude of downstream tasks, such as protein sequence design, protein folding modeling or protein-protein interaction prediction. In addition, we curate a feature-rich benchmarking dataset based on the latest annual release of the PDB and a selection of preprocessing parameters that are widely used across the research community. We showcase its utility by benchmarking a state-of-the-art (SOTA) deep learning model for protein sequence design. The open source code is packaged as a python library and can be accessed on https://github.com/adaptyvbio/ProteinFlow.

## 1 INTRODUCTION

The last decade has seen tremendous progress in the field of artificial intelligence thanks to the popularization of deep learning methods. Deep learning (DL) algorithms rely on neural networks, a computational model first proposed in the 1940s, in which layers of neuron-like nodes mimic how human brains analyze information [7]. Deep learning has achieved state-of-the-art performance in many applications, including image classification [65], object detection [39], machine translation [51], and voice recognition [16]. In recent years, deep learning approaches have been steadily infiltrating the realm of biology and biotechnology, with the promise of solving some of the great challenges of healthcare and biological industry. From drug discovery [34, 45, 67] to single-cell analysis [25, 53, 54] and genome editing [12, 59, 63], DL has demonstrated its value in enabling new discoveries in a field that has historically lagged behind the digital revolution [6].

One particular field that has seen an accelerated expansion of deep learning methods and techniques is protein design or engineering. DL has been successfully applied in many aspects of protein engineering, from sequence design [13, 49, 62] to function prediction [15, 24, 57]. One of the reasons for the great interest in applying machine learning to proteins is the diversity of functions carried out by proteins compared to other biological polymers [66]. Proteins are composed of only 20 different amino acid residues, yet they are involved in a wide range of biological processes, such as structural support, transportation, and regulation of gene expression [10]. Moreover, their local flexibility to selectively bind and specifically recognize other molecules makes them highly suitable for designing new therapeutic modalities [43]. Although all proteins are based on a limited set of building blocks, their combinatorial design space is vast and finding functional protein designs remains an unsolved challenge [50], hence the interest from the research community to apply DL algorithms to detect emerging trends that classical physics-based approaches cannot capture [41].

A multitude of new model designs and architectures have been proposed to improve the learning of protein representations, including attention strategies that are tailored to the invariances and symmetries of proteins [21], graph-based representations [13, 19], and model recycling strategies [2, 21]. However, as in other machine learning domains, data plays a crucial role, and research endeavors that focus on gathering and annotating biological data have been instrumental in moving the field forward.

Despite the numerous data modalities utilized for DL, protein macromolecular structures obtained through experimental methods such as cryogenic electron microscopy (Cryo-EM) or X-ray crystallography, are among the most valuable sources of protein data available. The biological function of a protein is directly linked to its resulting 3D structure, and hence this data provides crucial insights into how a protein is folded and how its different parts interact with each other and with other molecules [44]. Furthermore, 3D structure of a protein can explain how amino acid mutations can affect the catalytic activity [27], ligand-binding properties [56] or protein dynamic properties [28]. Given the richness and complexity of protein structural data, many deep learning approaches currently leverage this data for a multitude of protein engineering tasks [9]. However, due to the complexity of protein structural data, the task of filtering and processing the data for the different applications is not trivial and remains a barrier of entry for many researchers.

In general, labeling, classifying and representing protein structure data is complicated due to its hierarchical nature and high dimensionality [42]. Functional proteins are often composed of multiple chains, each of them made from hundreds or sometimes thousands of amino acids and each amino acid can be formed from tens of atoms. Every level of the protein hierarchy has unique features and characteristics that define the final properties of the protein ensemble, and capturing and aggregating all of these dependencies is challenging. Furthermore, selecting and filtering the correct protein data is a complicated task. For instance, large protein databases like the Protein Data Bank (PDB) [8], provide a vast set of macromolecular resolved structures in their native state, however the method and experimental conditions found for submitted structures are often heterogeneous and data can suffer from a range of issues including low accuracy structures, missing residues or redundancies [46, 48]. Furthermore, these datasets are heavily biased towards a few over-represented families of proteins, which can influence the downstream training tasks, if the data are not clustered correctly [37].

Another common problem is the lack of standardized training datasets in the field to support the growing number of protein modeling tasks. Several works have focused on processing the data contained in the PDB for DL applications [3, 31, 55]. However, these tools often suffer from several limitations. Firstly, datasets are designed for a specific protein modeling task (eg. protein structure prediction), and hence reusing and adapting the datasets to novel problems can be challenging and often requires reformatting or recomputing new features[31]. Most frameworks lack of convenient machine readable data formats for downstream analysis and often these datasets become outdated and do not reflect all the up-to-date data available. The latter is a great problem since protein structural data available today is quite limited and the PDB is constantly being updated with new structures -around 20′000 new structures are added annually, 10% of the total size of the dataset [33]. This lack of up-to-date data can significantly impede the downstream applications of deep learning models.

These issues are further exacerbated by the lack of a consensus in the research field for the selection of filtering criteria. There are a plethora of experimental parameters across the PDB, including the experimental method used to determine the structure or the resolution of the structure. However, selecting the correct parameters for a specific modeling task can be challenging [1]. Although there are certain rules-of-thumb in the field, there is currently no standard list of methods that are used across training datasets and there is limited research focus on how different data selection strategies can impact model accuracy [3]. This can lead to discrepancies in the data between groups and, consequently, a lack of reliable insight into the properties of the DL model.

Several initiatives have tried to provide a framework for testing and benchmarking models [11, 26, 31, 32, 38, 40, 52, 55, 58]. One of the best known initiatives in the protein structure modeling field is the Critical Assessment of protein Structure Prediction (CASP) [32]. CASP provides a blind assessment of protein structure prediction methods, which can be used to identify the strengths and limitations of the various models driving the development of new and improved computational tools. However, the training data used to benchmark the models is often not standardized and testing sets are small and incomplete. Furthermore, due to the rapid development of the machine learning field, this benchmarking sets are static or updated infrequently, preventing new models, tasks and modalities to be efficiently tested [3]. Another common problem is the lack of cohesion between the training and test sets, which often leads to information leakage across splits. Unlike CASP where the proteins in which the benchmarking set is hand-picked by experts in order to have varying levels of difficulty and ensure low homology overlap with the remaining protein structures available, most benchmarking datasets in the field do not follow this practice strictly, and its common to find similar protein structures across the training, validation and tests splits [11]. Hence it is important that standardized datasets are designed to accommodate a variety of training tasks, with a sufficient amount of data for training purposes and good safeguards against data leakage between train and test data.

Here we present a python package to exploit the vast amount of data found in the protein structure database. First, we provide a fully customizable end-to-end bioinformatic pipeline to extract, filter, annotate and cluster data from the PDB, allowing users to design and generate a custom dataset according to their specific modeling task and providing a ready-to-use output compatible with most machine learning frameworks. The pipeline allows to extract all levels of protein organization, including primary sequence, secondary structure, structural single chain information and protein complex information. At the end of the pipeline, pre-processed data are clustered and splitted in training, validation and test sets ensuring the absence of data leakage between the sets.

Secondly, we generate a ready to use clustered dataset for protein deep learning applications by using the most commonly used parameters and filters across the research community. We provide some insights on how these filters can affect model performances. We further validate the dataset by training a deep learning model for protein sequence design.

## 2 OVERVIEW

### 2.1 Dataset Processing Pipeline

In this section, we describe the methods used to produce a custom protein structure dataset using the *ProteinFlow* package, as well as a detailed explanation of all the supported features and parameters for filtering and clustering (Figure 1). The pipeline is packaged as a *python* library and is available at https://github.com/adaptyvbio/ ProteinFlow.

**Figure 1:**
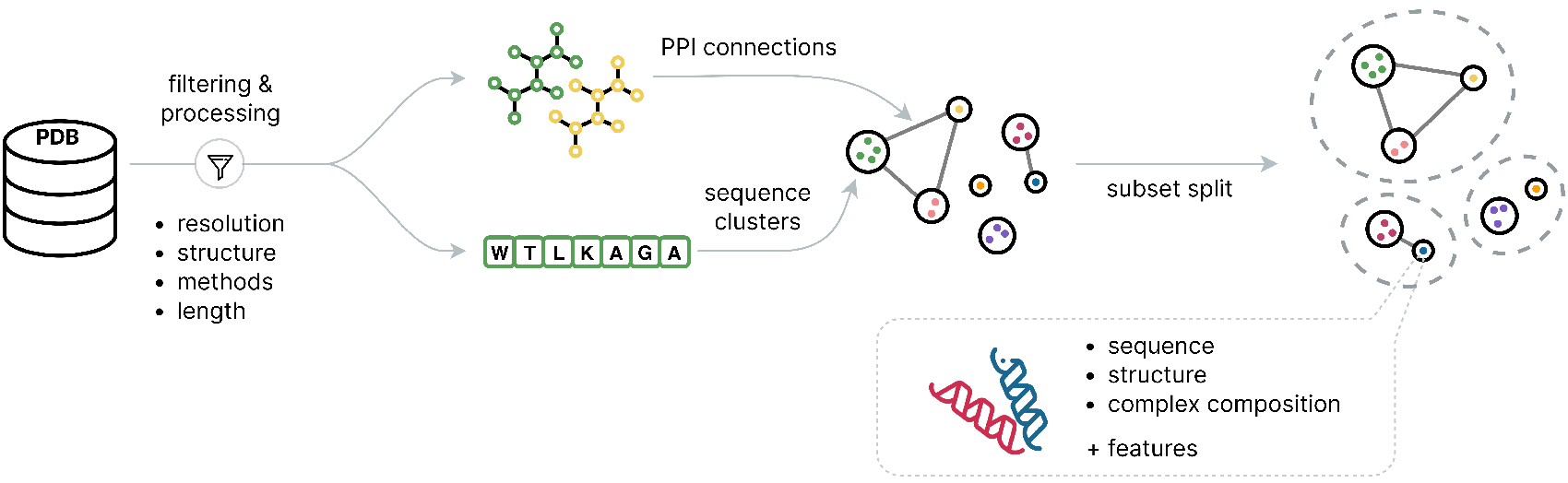
An overview of the pipeline. Biounit files are filtered and processed to extract aligned sequence and structure objects. Then sequence similarity clusters are linked in a graph according to protein-protein interactions. This graph is finally split into training, validation and test subsets.

### 2.2 Protein Organization in the PDB

In order to aggregate all levels of protein structure information it is necessary to understand how the protein organization is captured across experimental protein methods. Primary structure is represented by the sequence of amino acids in a polypeptide chain. This information is found in the PDB database in two formats: within the *PDBX/mmcif* files (excluding the residues with unresolved co-ordinates) and in associated .*fasta* files. The secondary structure of proteins refers to the local three-dimensional arrangement of the protein backbone atoms, specifically the pattern of hydrogen bonding between the amino acid residues [48]. Although this information is not explicitly annotated on the *PDBx/mmCIF*, there are several methods to compute it from the atomic coordinates [22]. Our package provides a method to calculate it from the pre-processed files. The tertiary structure describes the three-dimensional shape of a protein chain, which is determined by the spatial arrangement of its secondary structure, and it is explicitly found in *PDBx/mmCIF*. Finally, quaternary structure is the composition of interacting protein complexes. In the PDB this information is stored as biological assemblies also known as biological units (Biounits), which represent the macromolecular assembly that has either been shown to be or is believed to be the functional form of the molecule [65]. This is essential because using data that has not been functionally validated about protein interactions can lead to incorrect assumptions about the protein’s biological function, and the resulting models may not accurately represent the actual structure and interactions in vivo. For this reason, our pipeline processes and extracts all information from PDB biounits and in order to accommodate different modeling strategies we provide the option to keep the complexes intact, split them up into single chains or extract pairs.

#### 2.2.1 Download

For the stable versions of the dataset, we use the annual snapshot releases from the RCSB PDB [8]. Older released versions can be selected and used by the pipeline to create a different, more restricted dataset. Accessing all the up-to-date entries from the PDB dataset can be done by using the date filtering as outlined below. For efficiency, files are not extracted directly from the PDB website or the PDB archive but from the *Registry of Open Data on Amazon Web Services (AWS)* [5]. The PDB offers a variety of file formats for macromolecular structure records. The most common ones being the *PDB* and the *PDBx/mmCIF* file formats. The pipeline uses by default the *PDBx/mmCIF* file format as it has become the standard file for structure deposition for the wwPDB database since 2014 and also this format has no size restrictions compared to the older *PDB* format, hence being able to access even large protein assemblies [60].

#### 2.2.2 Filtering

The pipeline involves two levels of filtering. The initial level of filtering is done by querying directly the PDB using the PDBe REST API [33]. The PDBe API allows up-to-date programmatic access to the PDB database and supports several high level queries that can be used to search and select desired protein characteristics. The first filtering step outputs a list of PDB IDs, which are then downloaded as previously described. On the second filtering step, the downloaded *PDBx/mmCIF* files are processed, to extract all the relevant biological data (ex. coordinates, residues, chains, etc). At this step we also apply several data quality filters to ensure that files with erroneous or missing data are not integrated into the final dataset. All filters can be adjusted or modified unless stated otherwise and their usage can be found in the package documentation.

- **Method:** The experimental methodology used to determine the protein structure (eg. X-ray crystallography, NMR spectroscopy, etc). For the stable release, we select entries that were resolved using X-ray crystallography or electron microscopy and reject other experimentally determined methods such as Nuclear Magnetic Resonance (NMR). This is a commonly extended practice across training datasets in the field [3, 21, 36]. In general there are two reason for this filtering decision. First, the number of structures available from NMR or other experimental methods is very small compared to the selected methods -13,000 structures as of Feb 2023 - and second, NMR structures can have larger uncertainties than X-ray or cryo-EM structures, particularly in regions with high flexibility or low structural density [17], making them less suitable for training deep learning models based on protein structure. On the other hand, there are some deep learning approaches that have been developed to predict protein structures from NMR data, including several recent studies [23, 30].
- **Resolution:** The threshold on the refinement resolution above which we discard an entry. This filter is used to remove any low resolution entry, which can impact data quality. This value oscillates between 0.48 and 70 Angstroms in the PDB as of today. For the stable release, we select entries with a resolution below 3.5 Angstroms. The resolution threshold for experimentally determined structures used across the field varies depending on the method and the specific study. In general, higher resolution structures (i.e., those with lower values of the crystallographic R-factor or the cryo-EM map resolution) tend to be a better source of data for protein modeling tasks and the resolution threshold is commonly accepted to be between 0.5 and 4.0 Angstroms [3, 21, 64]. It is typically independent of the protein modeling tasks. There have been a few studies that have tested the effect of training on lower quality structures on model performance [35]. These studies suggest that while higher-quality structures are generally preferred for training deep learning models, lower-quality structures can still be useful for training and testing, and can help improve the robustness and generalizability of the models.
- **Polymer entity:** The types of polymers that are allowed in the biounits. This filter cannot be changed and it selects entries that only contain protein and oligosaccharide chains. The current version of the *Protein Data Bank* (PDB) contains around 200′000 resolved structures of biological entities, most of which are proteins [8]. Other types of entities that can commonly be found in the PDB are nucleic acids (ex. DNA), carbohydrates, glycans, lipids and small molecules. Although these entities play fundamental regulatory and functional roles in organisms and can be of interest for some specific modeling applications such as protein-ligand interaction prediction, they are often not considered or represented in general protein modeling tasks, such as protein structure prediction or protein-protein interaction. Furthermore, the presence of this entities can greatly affect the conformation of proteins and induce biases. The next set of filters can only be applied by accessing the information inside the *PDBX/mmCIF* and are thus performed after downloading the previously filtered entities. Apart from filtering the dataset for specific requirements, at this step we also apply a set of quality control methods to remove incomplete or mislabeled data and ensure data robustness across all datasets.
- **Sequence size:** Some proteins are actually very small and are thus quite flexible, much like a small peptide. Taking them into account to predict structure is likely to hamper any learning or testing procedure since the resolved structure could be one of many. For the stable release, we choose a cutoff threshold of 30 residues (after removing missing residues) under which proteins are discarded. As before, biounits are selected only if all the chains they contain satisfy this filter.
- **Redundancies:** Some biounits coming from a same PDB submission are actually identical sequence-wise. This is due to the spatial repetition of structures in a crystal and thus these biounits represent the same structure(s). Sometimes, some chains can be missing in one of these similar biounits because they have not been resolved. To eliminate redundancies, we opted to retain only one biounit among those that originate from the same PDB and share identical sequences (defined as at least 90% sequence identity for greater robustness). Additionally, we discarded all biounits whose sequences are entirely contained within another biounit’s sequence set from the same PDB.
- **Missing residues or atoms:** A residue is considered as missing if any of its backbone atoms are missing. When a large set of residues are missing, recomputing the location of the 3D coordinates is difficult and prevents the sample to be used for training or testing purposes. Residues in the middle of the protein chains are crucial to define the protein 3D structure as they often form part of the stabilizing protein core [48], while information close to the tails often interacts less strongly with the rest of the protein. Furthermore, during sample preparation and purification it is a common practices to add short affinity tags to the ends of proteins, which often are present and in principle should not affect substantially the the protein’s fold [14]. The filter is designed to distinguish between the different locations of missing residues and both thresholds can be adjusted independently. For the stable release, we only select proteins that have less than 30% missing residues in the tails and less than 10% missing residues in the middle (sequence with missing residues in the tails removed). Biounits are then selected if all the chains they contain satisfy this filter.
- **Unnatural amino acids:** Some proteins contain unnatural amino acids. Although, several studies have focused on modeling the structure and interactions of novel residues in proteins [47], in general the presence of non-canonical amino acids in the PDB is rare and most recent models do not integrate this information during learning [21]. The pipeline is designed to discard every biounit that contains unnatural amino acids. This filter cannot be changed.
- **Unexpected atoms:** Each amino acid is composed of a specific set of atoms. Side chain atoms are commonly missing, but in rare occasions additional atoms can be found under the same residue. We check that the residues only contain atoms they are supposed to contain and discard all proteins (and corresponding biounits) that contain any additional atoms. This filter cannot be changed.
- **Discrepancies between fasta and PDB sequences:** During the processing of the biounits, sequences from the fasta (complete sequences) are aligned to the sequences extracted from the PDB. This allows to identify the positions of the missing values as well as link amino acid features to their right position in the sequence. In some rare cases, alignments do not match or some chains in the PDB do not appear in the fasta, and the concerned biounits are discarded. This filter cannot be changed.
- **Large biounits:** Some biounits are particularly large, containing hundreds of chains, for example for structures of a virus capside or a chaperone.

In the stable release, we discard all biounits that contain chains with more than 10^′^000 amino acids in total (including missing values).

#### 2.2.3 Clustering and splitting

Clustering can be done either at the sequence or at the structure level. The CATH database provides structural clusters of protein domains, and several works have used these CATH topology classes to cluster and split their dataset [4, 18, 20]. However, CATH is not a tool that can output a classification on any dataset it is given. Rather, it is a classification that is regularly updated but that may lack the up-to-date set of entries submitted to the PDB.

Therefore, we perform clustering based on sequence identity, which yields equivalent results, but can be used on any dataset without losing any entry. To this end, we use the *MMseqs2* suite [29] and cluster proteins at 30% sequence identity, as in [13] (this parameter can be changed).

Our goal is to cluster biounits, not protein chains. Indeed, if the dataset is used in the context of protein-protein interactions, we need to take care that there is no information leakage caused by a biounit containing both a chain in the *train* set and one in the *validation* set for example.

In order to go from sequence clustering to biounit clustering, we build a graph where nodes are the clusters produced by *MMseqs2* and edges between two nodes represent the number of chains in the two nodes that come from the same PDB as a chain in the other node. We call this a protein-protein interaction (PPI) graph. See the statistics for the 2022 stable release version of this graph on Figure 2 A-C. Each connected component of this graph is then defined as a biounit cluster since no biounit is shared between two of these connected components.

**Figure 2:**
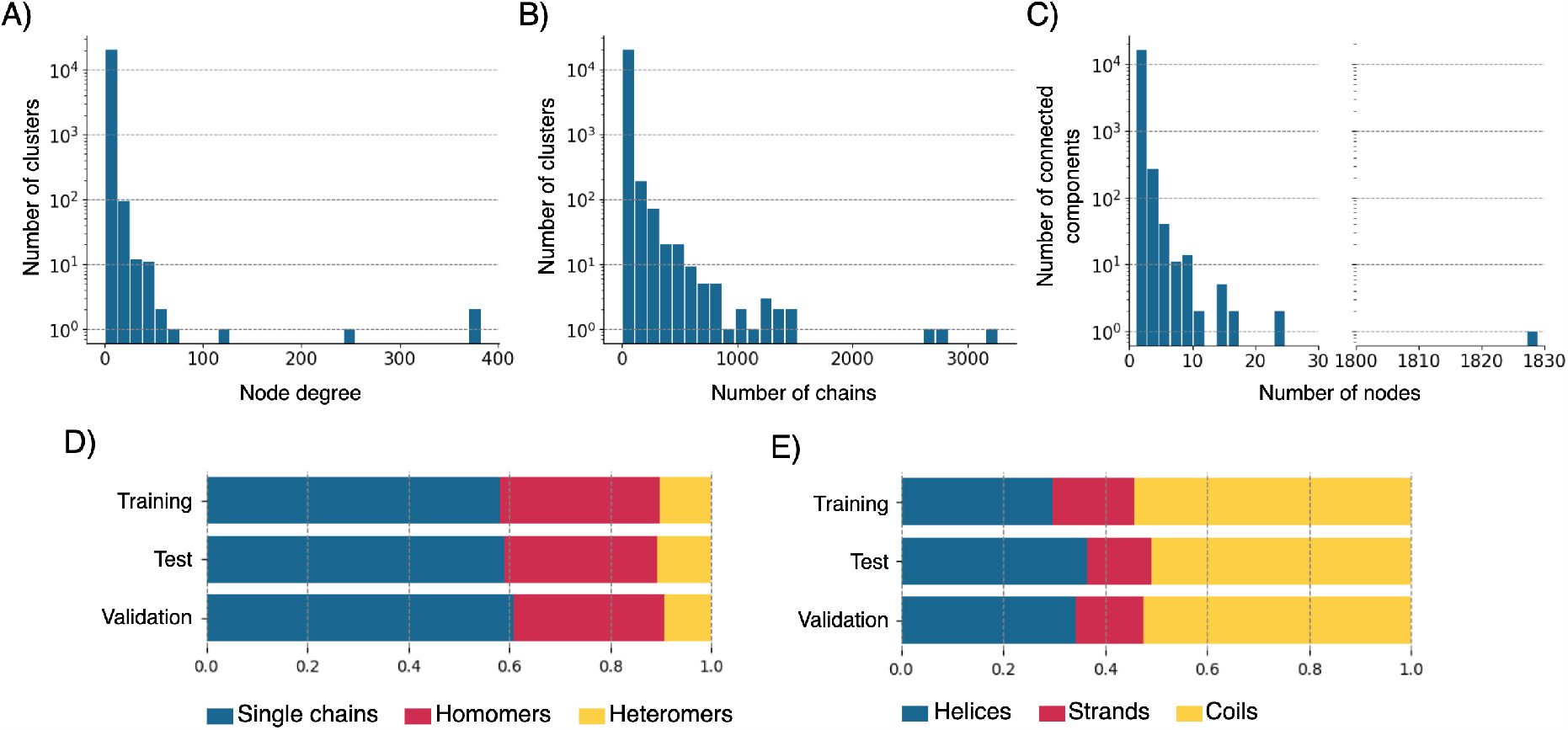
Statistics for the 2022 stable release. Histograms for A) the node degree in the PPI graph and B) the number of chains over MMseqs clusters. C) The distribution of size across connected components in the PPI graph. Distributions of D) single chains, heteromers and homomers and E) secondary structure elements in the training, validation and test subsets.

For any deep learning application, we need to split the dataset into *train, validation* and *test* sets. A random splitting of the clusters according to the split ratios is in theory possible to achieve this purpose. However, we have here a heterogeneous dataset, in the sense that it contains biounits with only a single chain alongside with biounits with a homomer and biounits with a heteromer. As these different subtypes are likely to follow slightly different rules (the amino acid composition of a binding site differs from the composition of the free surface, and it is also dependant on the content of the interacting chains [48], we want to take care that the distributions of *single chains, homomers* and *heteromers* are similar between the *train, validation* and *test* sets.

**Table 1:** Number of biounits removed for each filter in the second filtering stage for 2022 ProteinFlow Stable dataset. The data shows that a large number of entities within the PDB contained several issues that limit the exploitation of the data for deep learning purposes.

| Filter | Number of biounits |
| --- | --- |
| Redundancies | 63'418 |
| Missing values | 22'382 |
| Sequence size | 9'824 |
| Unnatural amino acids | 626 |
| Unexpected atoms | 146 |
| Discrepancies between fasta and PDB sequences | 181 |
| Missing sequence files | 7 |
| Large biounits | 15326 |

Consequently, when creating the dataset, we also split it while taking this constraint into account. In practice, we first try to partition the dataset randomly. If after fifty trials the partition still does not meet the criteria (total number of sequences inside each set as well as similar distributions of *single chains, homomers* and *heteromers*) within a tolerance margin (here 20% of the split ratios), we add and remove clusters one by one until we reach the criteria. The tolerance is needed since we are dealing with clusters of sequences and the ratios apply on the number of sequences.

It should be noted that the splitting algorithm is tailored for big datasets and that it is not guaranteed to yield a good re-partition for very small datasets (around 250 biounits or under). The algorithm is designed so that if one of the three classes (*single chains, homomers* or *heteromers*) is highly underrepresented compared to the size of the dataset (which is almost certain to happen in the case of small datasets), the re-partition criteria between the sets are ignored for this specific class. To handle such edge case is useful to conduct tests but the results should not be trusted in another context.

While building the dataset, we noticed that the biggest biounit cluster for the 2022 stable release contained 56^′^430 chains, which represents around 20% of the whole dataset (see a visualisation of this component on Figure 3). In the 2022 PPI graph, it contains 84 out of the 100 nodes with the largest node degrees. Given its size, the cluster is always assigned to the *training* set. The distribution of cluster sizes can be seen on Figure 2 C.

**Figure 3:**
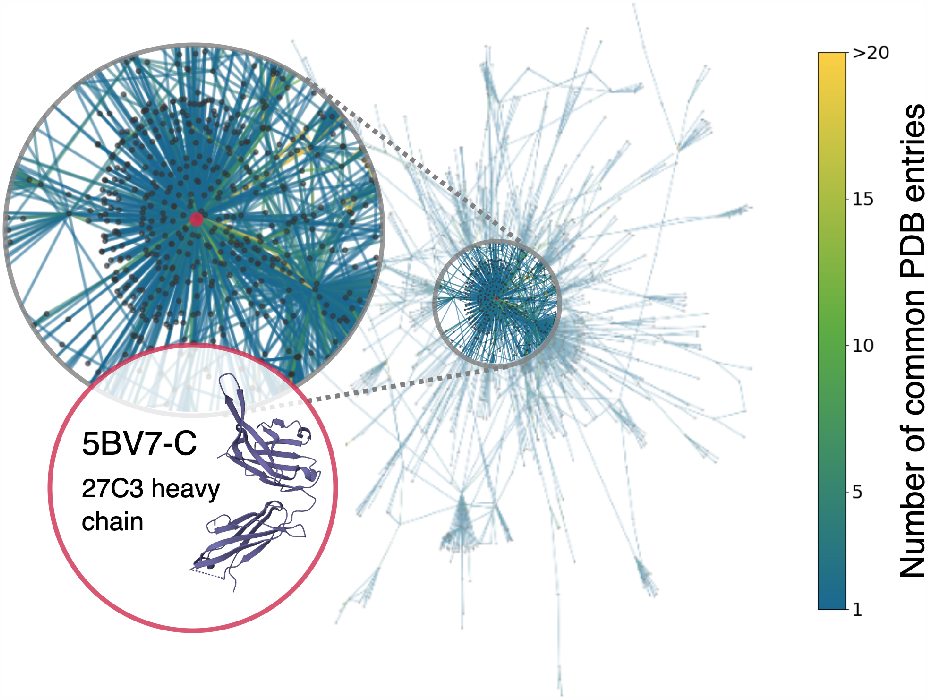
The largest connected component of the PPI graph generated using the MMseqs2 package. The red dot represents the cluster with the largest node degree, i.e largest number of . This cluster contains antibody heavy chains and homologs as shown by one of the cluster representatives, the 27C3 heavy chain.

The apparition of such huge clusters is actually due to the fact that several protein families are over-represented on the PDB. For instance, antibodies are a class of proteins that due to their therapeutic implications and their great designability properties, they have been extensively studied. Furthermore, antibodies can selectively bind to many protein targets, while sharing a high sequence similarity with other members of the family [61]. Most of the nodes with the largest degrees indeed represent antibody chains. For instance, the representatives of the 3 nodes with the largest degrees are 27C3 heavy chain (PDB ID: 5BV7-C), 3C10 Fab’ light chain (PDB ID: 5W5X-L) and iv8 light chain (PDB ID: 6OL7-I).

#### 2.2.4 Data format

Data contained within the curated dataset is split into four subfolders, in order to leverage the data for different downstream tasks:

1. **splits_dict**: contains the dictionaries with the clusters information
2. **training**: contains all the biounits present in the *train* set (*pickle* format)
3. **validation**: contains all the biounits present in the *validation* set (*pickle* format)
4. **test**: contains all the biounits present in the *test* set (*pickle* format)

The *split_dict* subfolder contains three *pickle* files: *train*.*pickle, valid*.*pickle, test*.*pickle*. Each file contains two python dictionaries. The first one is organized by cluster: each key is the name of a biounits cluster and the corresponding value is a *numpy array* that lists all the chains in the cluster in the format *[biounit_file_name, chain_name]*. The second dictionary contains the same information with an additional hierarchy layer on top by type of chain (*single chains, homomers* or *heteromers*). This way one can only select one type of chain depending on the desired application. For instance, if one wishes to access the biounits in cluster *1d7b_A-B*, which is a cluster of homomers, it can be done so by writing `dict[‘homomers’][‘1d7b_A-B’]`.

A `Dataset` class is provided to load the data in *PyTorch*. It is designed to sample elements from the clusters and aggregate them into a batch, with the possibility to compute some features that could be needed as input to a model. The current supported features are: secondary structure, dihedral angles, chemical properties and sidechains orientation in the global frame. Chemical properties are represented by a vector of six features: residue hydropathy, volume, charge, polarity and electronic behavior (electron acceptor or donor). Sidechain orientation are defined as the vector originating from the residue *C*_*α*_ and ending in one predefined heavy atom in the sidechain, that is chosen for each amino acid type so that the resulting vector is representative of the global orientation of the sidechain. All vectors are normalized. In addition, this class supports sampling a different random chain from the sequence identity clusters at every iteration, as in [13]. This way, it is possible to avoid biasing a model towards more widespread protein families while still leveraging the power of a large dataset.

**Table 2:**
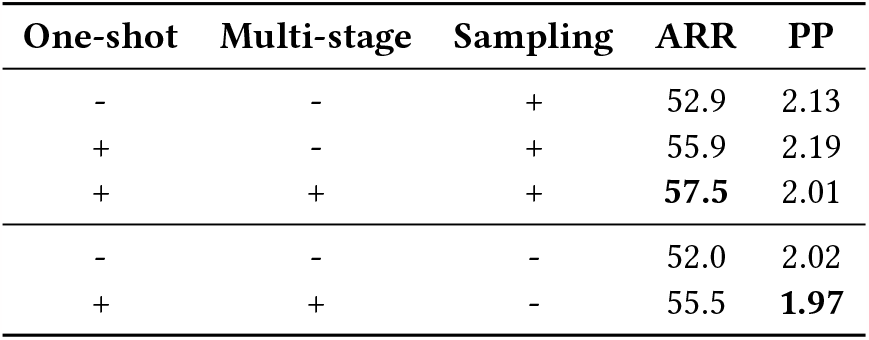
Amino acid recovery rates and perplexities with different model modifications.

## 3 BENCHMARKS

### 3.1 Dataset statistics

While avoiding data leakage is very important in defining the train/validation/test split for a benchmark, it is not the only criterion for a good partition. We also need to make sure that the validation and test subsets provide a good representation of the general data distribution in order to avoid introducing bias in model selection. In order to check for this, we compare the the proportions of single chains, homomers and heteromers (Figure 2 D), as well as distributions of secondary structure elements across the subsets (Figure 2 E).

### 3.2 Sequence design study

As a case study, we used our dataset to test how the performance of a sequence design model (ProteinMPNN, [13]) changes with the introduction of several architecture modifications. In particular, the task was to reconstruct the sequence of whole chains given the structure and the sequence of other chains in the biounit, if they exist. The results can be found in **??**.

We add the following changes to the model:

- **One-shot prediction:** The original model outputs sequence prediction in autoregressive mode. That means that the model iterates over the amino acids in a sequence, generating the probability distribution for every new residue given the previous predictions. We suggest an alternative approach: generating all predictions in one operation (*One-shot* in the table).
- **Multiple stages:** We increase the number of parameters in the model by introducing multiple stages of encoders and decoders. Before the first stage, the sequence information is masked, and before every other starting from the second it is initialized with the predictions from the previous stage (*Multi-stage* in the table).

In addition, we assessed the influence of random sampling on the performance of the model (*Sampling* in the table). In the original paper, the multi-chain ProteinMPNN model was trained with choosing a new random representative for each sequence similarity cluster at each pass through the dataset [13]. This mode of training is also available in proteinflow and our experiments show that it can improve the results. The code for training the models is available on GitHub.

## 4 LIMITATIONS AND FUTURE WORK

As of now, ProteinFlow is limited to the study of proteins, excluding other important polymers such as DNA or small ligands. It would thus be interesting to add the possibility to handle such entities in the pipeline, given that they are non-negligible part of biological interactions in a cell. Additionally, we would like to add some important features such as residue contact features to further generalize the use-cases of this pipeline.

## 5 CONCLUSION

Here we present a flexible data processing pipeline, as well as a benchmarking dataset for protein design. We hope that it will contribute to the advancement of deep learning research in protein design by providing a ground for method comparison and the means for easier data generation and customization across the growing field.

